# MicroED Structures from Micrometer Thick Protein Crystals

**DOI:** 10.1101/152504

**Authors:** Michael W. Martynowycz, Calina Glynn, Jennifer Miao, M. Jason de la Cruz, Johan Hattne, Dan Shi, Duilio Cascio, Jose Rodriguez, Tamir Gonen

## Abstract

Theoretical calculations suggest that crystals exceeding 100 nm thickness are excluded by dynamical scattering from successful structure determination using microcrystal electron diffraction (MicroED). These calculations are at odds with experimental results where MicroED structures have been determined from significantly thicker crystals. Here we systematically evaluate the influence of thickness on the accuracy of MicroED intensities and the ability to determine structures from protein crystals one micrometer thick. To do so, we compare *ab initio* structures of a human prion protein segment determined from thin crystals to those determined from crystals up to one micrometer thick. We also compare molecular replacement solutions from crystals of varying thickness for a larger globular protein, proteinase K. Our results indicate that structures can be reliably determined from crystals at least an order of magnitude thicker than previously suggested by simulation, opening the possibility for an even broader range of MicroED experiments.

**Summary:** Atomic resolution protein structures can be determined by MicroED from crystals that surpass the theoretical maximum thickness limit by an order of magnitude.

## Main Text

### Introduction

Electrons interact with matter more strongly than X-rays and offer a larger fraction of useful, elastic scattering events to inelastic scattering events (1). These properties are leveraged by the cryoEM method, MicroED, for structure determination at atomic resolution from nanoscale protein crystals (2). With this method, diffraction is measured from protein nanocrystals in a frozen-hydrated state (3) using a low dose electron beam (typically ~0.01 e^-^ Å^-2^ s^-1^) (4–6). Crystals are continuously and unidirectionally rotated in the beam while diffraction images are collected as a movie on a fast detector (7). A number of previously unknown as well as known structures have been determined by this method to atomic resolution (8).

The strong interaction between electrons and materials also allows for multiple scattering events to take place before electrons exit the specimen (9). According to dynamical scattering theory (10), multiple scattering events produce inaccuracies in the recorded reflections, potentially preventing the solution of structures. Simulations suggest that with crystals thicker than 50-100 nm (11, 12) dynamical scattering can be severe, resulting in nearly random intensities, where the relationship between the intensity and structure factor no longer holds true (10). However, these simulations assume diffraction is recorded from a perfect and stationary crystal - real macromolecular crystals are not perfect. In fact, precession electron diffraction can avoid many of the artifacts associated with dynamical scattering from near perfect crystals of inorganic material by pivoting of the electron beam around the crystal (13). Similarly, by employing continuous rotation MicroED (7), useful diffraction is routinely collected from protein crystals hundreds of nanometers thick (14). Even 1.5 μm-thick crystals of lysozyme were shown to produce diffraction that when integrated produced reasonable statistics (2). Moreover, recent structures determined by *ab initio* methods to 1Å resolution further indicate that the diffraction intensities obtained by continuous rotation MicroED are accurate and maintain the relationship between amplitude and phase (4, 15).

Here we systematically investigate the relationship between crystal thickness, dynamical scattering and the quality of structure solutions obtained by MicroED. We determine *ab initio* structures of a segment from the β_2_-α_2_ loop of human prion protein in the amyloid state as well as the structure of proteinase K by molecular replacement from crystals up to a micrometer thick.

### Results

#### A comparison of structures determined from thin and thick crystals of a segment of the β2-α2 loop of human prion protein

As a model system for evaluating the effects of crystal thickness on diffraction intensities, we use a segment from the β2-α2 loop of human prion protein (hPrP) that contains a glycine residue at its amino terminus (sequence GSNQNNF), hereafter referred to as hPrP-β2α2, for MicroED structure analysis from crystals that vary in thickness. The crystals of this segment appear as micrometer-long needles and vary in thickness from several nanometers to over a micrometer. Seven data sets originating from thin crystals were collected and combined to yield a reference data set that was 80.3% complete in P1 with constants {**a**, **b**, **c**} (Å) = {4.86, 14.11, 18.41}, and angles {α, β, γ} (°) = {90.00, 93.71, 101.21}. A second data set was constructed from 7 thick crystals between 500 nanometers and one micrometer thick, that were combined to yield a 75.6% complete data set with the same space group and unit cell dimensions as above. Thin and thick crystal structures were determined *ab initio* by direct methods using SHELXT (16) (See **SI Methods**) and refined to atomic resolution using BUSTER-TNT (17) showing clear atomic density (**Figure 1)**. Refinement statistics for hPrP-β2α2 are presented in **Table 1**.

**Figure-1:**
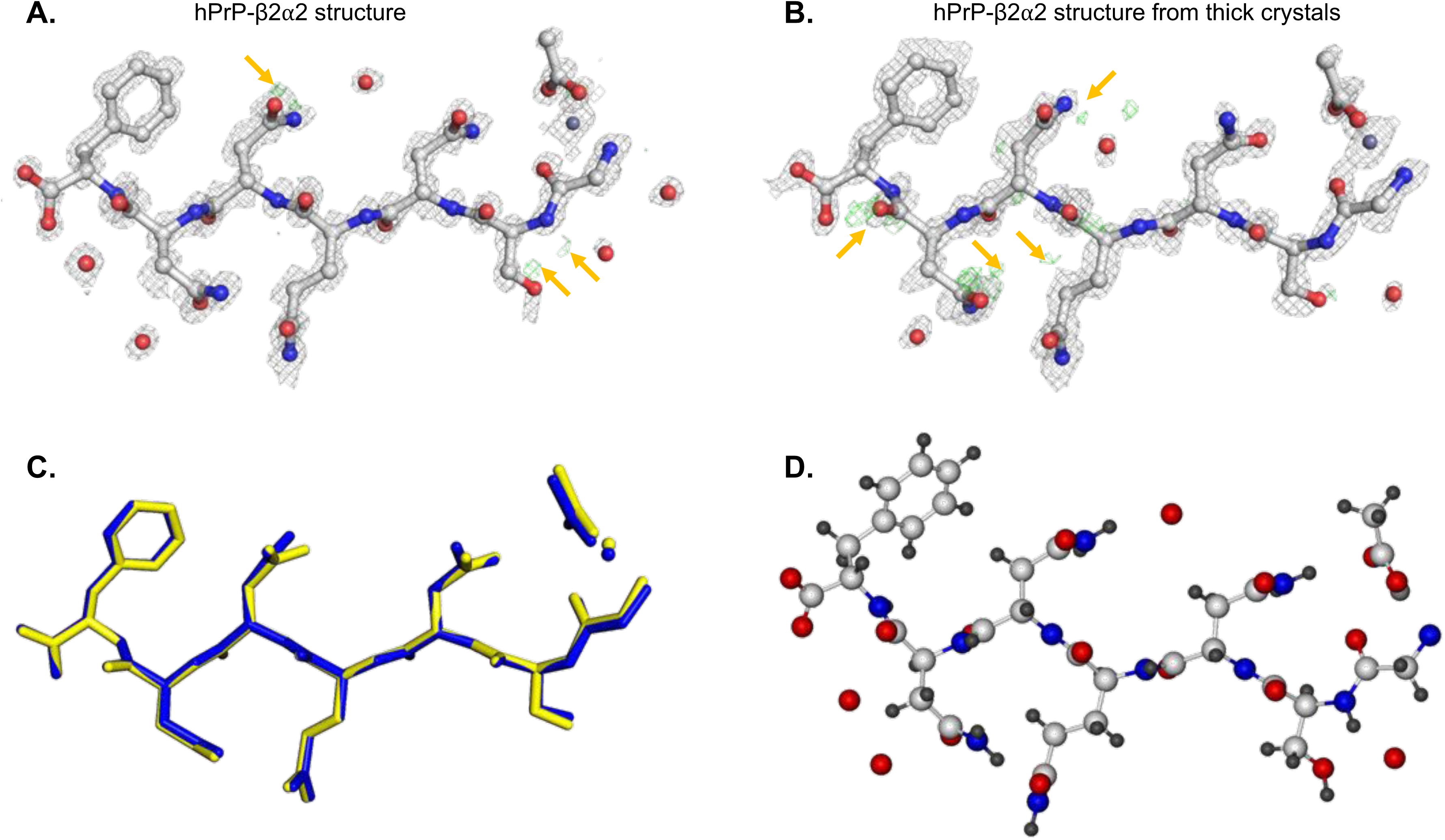
Structural comparison of hPrP-β2α2 from thin and thick crystals by direct methods. (**A**) Structural model for hPrP-β2α2 solved from typically <200 nm crystals (**B**) from a set of crystals with average thicknesses >1 μm. 2mFo-DFc and mFoDFc density maps are contoured at 1.5 σ and 3 σ levels, respectively. Positive hydrogen densities for both solutions are visible in **A** and **B** in green and are designed by orange arrows. (**C**) *Ab initio* solutions from both normal (yellow) and >1 μm (blue) peptide crystals. (**D**) Final structural model of hPrP-β2α2.

**Table-1:** Data Collection and Refinement Statistics.

|  | hPrP-β2α2 | Thick hPrP-β2α2 | Proteinase K | Proteinase K<br>(Intermediate Thickness) | Proteinase K<br>(>1μm, single crystal) | Proteinase K<br>(>1μm, multiple crystals) |
| --- | --- | --- | --- | --- | --- | --- |
| Wavelength (Å) | 0.0251 | 0.0251 | 0.0251 | 0.0251 | 0.0251 | 0.0251 |
| Resolution range (Å) | 7.608 - 1.001 (1.037 - 1.001) | 14.03 - 1.011 (1.047 - 1.011) | 19.62 - 1.75 (1.813 - 1.75) | 19.39 - 2.0 (2.072 - 2.0) | 22.16 - 3.001 (3.108 - 3.001) | 22.61 - 2.5 (2.589 - 2.5) |
| Space group (#) | P 1 (1) | P 1 (1) | P 43 21 2 (96) | P 43 21 2 (96) | P 43 21 2 (96) | P 43 21 2 (96) |
| Unit cell<br>a, b, c (Å) | 4.86 14.11 18.41 | 14.34 4.89 17.8 | 67.32 67.32 101.01 | 67.21 67.21 100.59 | 68.09 68.09 102.13 | 67.5 67.5 102.41 |
| α, β, γ (°) | 90 93.71 101.21 | 94.96 91.02 101.82 | 90 90 90 | 90 90 90 | 90 90 90 | 90 90 90 |
| Total reflections | 12907 (524) | 8654 (307) | 104808 (5429) | 57404 (4205) | 20317 (1505) | 78690 (5656) |
| Unique reflections | 2072 (171) | 1869 (141) | 22698 (2256) | 15756 (1543) | 4135 (294) | 8627 (828) |
| Multiplicity | 6.2 (4.2) | 4.6 (3.2) | 4.6 (4.4) | 3.6 (3.6) | 4.9 (5.1) | 9.1 (9.2) |
| Completeness (%) | 80.29 (69.08) | 75.90 (60.00) | 94.02 (94.14) | 97.18 (97.60) | 79.70 (81.82) | 99.41 (99.64) |
| Mean I/sigma(I) | 5.05 (2.25) | 4.64 (1.65) | 3.1 (1.0) | 3.0 (0.87) | 2.20 (0.92) | 4.83 (2.39) |
| Wilson B-factor | 4.38 | 6.55 | 11.77 | 24.23 | 39.66 | 17.86 |
| R-merge | 0.238 (0.43) | 0.233 (0.474) | 0.458 (1.989) | 0.294 (1.28) | 0.551 (1.47) | 0.414 (0.960) |
| R-meas | 0.257 (0.485) | 0.258 (0.552) | 0.516 (2.258) | 0.347 (1.51) | 0.617 (1.63) | 0.44 (0.97) |
| CC<sub>1/2</sub> | 0.98 (0.86) | 0.979 (0.489) | 0.929 (0.112) | 0.972 (0.289) | 0.883 (0.316) | 0.939 (0.484) |
| Reflections used in refinement | 2072 (172) | 1869 (141) | 22670 (2235) | 15756 (1544) | 4135 (414) | 8627 (828) |
| Reflections used for R-free | 207 (17) | 180 (12) | 1476 (146) | 788 (77) | 207 (20) | 431 (41) |
| R-work | 0.1984 (0.2067) | 0.1925 (0.2102) | 0.217 (0.348) | 0.2218 (0.3112) | 0.1987 (0.3059) | 0.2085 (0.2858) |
| R-free | 0.2382 (0.222) | 0.2523 (0.2920) | 0.266 (0.418) | 0.2508 (0.3390) | 0.2497 (0.4124) | 0.2612 (0.4195) |
| Number of non-hydrogen atoms | 66 | 63 | 2170 | 2130 | 2037 | 2090 |
| macromolecules | 55 | 55 | 2027 | 2027 | 2027 | 2027 |
| ligands | 5 | 5 | 10 | 10 | 10 | 10 |
| solvent | 6 | 3 | 133 | 93 | 0 | 53 |
| Protein residues | 7 | 7 | 281 | 281 | 281 | 281 |
| RMS (bonds) | 0.014 | 0.013 | 0.003 | 0.004 | 0.005 | 0.003 |
| RMS (angles) | 1.82 | 1.69 | 0.57 | 0.74 | 0.92 | 0.7 |
| Ramachandran favored (%) | 100 | 100 | 97.11 | 97.11 | 97.11 | 97.11 |
| Ramachandran allowed (%) | 0 | 0 | 2.53 | 2.53 | 2.53 | 2.53 |
| Ramachandran outliers (%) | 0 | 0 | 0.36 | 0.36 | 0.36 | 0.36 |
| Rotamer outliers (%) | 0 | 0 | 0 | 0 | 0.94 | 0 |
| Clashscore | 0 | 0 | 4.54 | 4.81 | 3.29 | 2.28 |
| Average B-factor | 5.85 | 6.78 | 11.49 | 23.19 | 25.12 | 12.61 |
| macromolecules | 4.76 | 6.05 | 11.19 | 23.28 | 25.15 | 12.75 |
| ligands | 7.48 | 9.69 | 29.54 | 27.28 | 19.83 | 16.75 |
| solvent | 14.52 | 15.18 | 14.63 | 20.75 | 0 | 6.39 |

The structure of hPrP-β2α2 represents a prion protofilament with amyloid features. The protofilament is a class 2 steric zipper, composed of parallel, face-to-back beta sheets (18). One pair of sheets makes the protofilament, as observed with other amyloid structures (15, 18). At this resolution, the density shows hydrogen atoms as well as the presence of zinc and acetate ions that facilitate crystallographic contacts; both were present in the crystallization condition (**Figure 1**). Structures from both thick and thin crystals show clear densities for waters in the 2F_o_-F_c_ map and have multiple hydrogens appearing in the F_o_-F_c_ density at the 3 σ level (**Figure 1**). The appearance of resolvable hydrogens in electron diffraction was first reported for proteins by Rodriguez et al. (15) and later by Palatinus (19) for small molecules. Our *ab initio* solutions from thin and thick crystals are very similar, with a backbone RMSD of 0.07 Å and an all-atom RMSD of 0.09 Å (**Figure 1)**. In summary, we found no significant differences between structures determined from thin and thick crystals of hPrP-β2α2.

#### A comparison of structures determined from thin and thick crystals of a globular protein

We collected MicroED data from single ~500 nm and ~1 μm-thick crystals of proteinase K (**SI Figure 2, SI Figure 3**) and determined structures from each. Reflections were recorded to a resolution of 1.8 Å in both cases yielding 97% completeness for the 500nm crystal and 79.7% completeness for the 1 μm thick crystal. Data were reduced in XDS (20) and phased by molecular replacement using the atomic coordinates of PDBID **5i9s** as a search model (21). Structures were refined using phenix.refine (22) with a high-resolution cutoff of 2.0 Å and 3.0 Å, respectively (**Table 1**; **Figure 2**). As the single, ~1 micrometer thick crystal structure had poor statistics, its data were merged with data gathered from three additional crystals of proteinase K; each of these also ~1 μm thick (**SI Figure 4, SI Figure 5**). The structure from the four combined data sets was determined again using the same search model now to a resolution of 2.5 Å with better overall statistics (**Table 1**, **Figure 2**). A comparison of the resulting structures with the molecular replacement search model indicate good agreement with lower than 0.25Å all atom RMSD. In summary, we found no significant differences between structures determined from thin and thick crystals of proteinase K.

**Figure-2:**
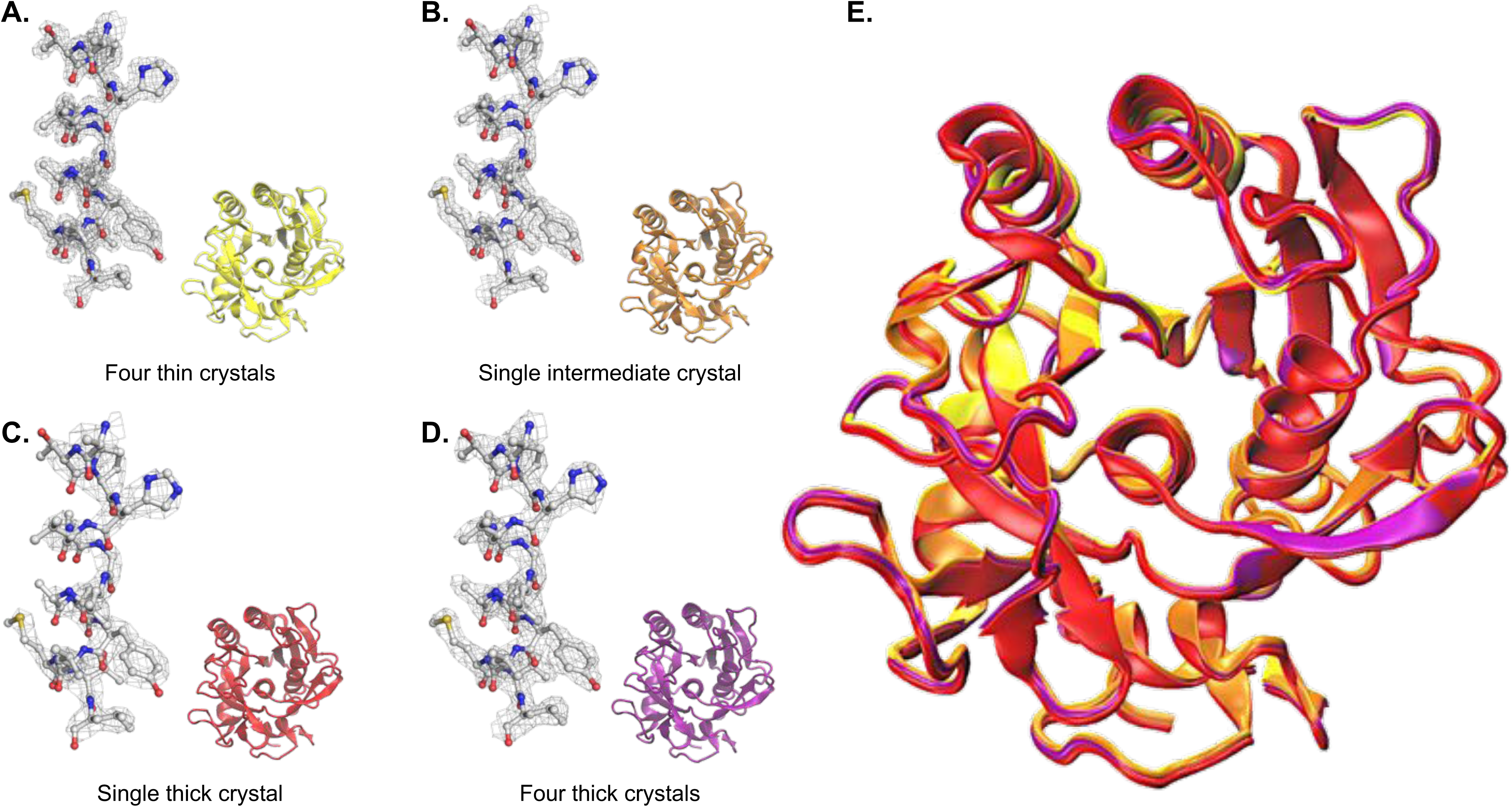
Structural comparison of globular proteins determined from thin, intermediate and thick crystals. Proteinase K sidechain density and corresponding structure solution from (**A**) thin ~200nm crystals (PDBID 5I9S)(20), (**B**) a single 600 nm-thick crystal, (**C**) a single 1000 nm-thick crystal, and (**D**) four crystals thicker than 1000 nm. All 2F_o_-F_c_ density maps are contoured at the 1.5σ level for residues 226-240 shown as a grey mesh. (**E**) All structure solutions of proteinase K overlaid for comparison. Overlaid colors correspond to individual colored structures.

#### Systematic study of the effects of crystal thickness on MicroED data quality

Diffraction was measured from 19 crystals of hPrP-β2α2 with thicknesses ranging from ~100 nm to ~1100 nm over similarly-sized wedges of reciprocal space corresponding to a real-space angular range of approximately -30° to +30° (**SI Table 1**). Data for all comparisons was indexed and integrated using XDS (20). Estimation of crystal thickness is discussed at length in **SI Materials**. Briefly, average crystal thickness is estimated by measuring the projected area from images recorded at 0° and 60° tilt, and the aspect ratio fit to an idealized model. These geometrical estimates were corroborated by intensity ratios using camera counts as previously described (14, 23) and are in good agreement (**Si Figure 6**). Crystal images are presented in **SI Document 2**. To assess the quality of diffraction data, we compare R values (Rmeas), the ratio of intensity to variance (I/σI), and the half-set correlation coefficient (CC1/2). All measured data values and statistics are presented in **SI Table 1**. Measured R values for these 19 crystals show a mean value of 13.03% with a standard deviation of 2.4%; average I/σI values average 4.1 with a standard deviation of 0.91. The half-set correlation coefficient is on average 98.2% with a standard deviation of 1.4%. The structure factor amplitudes for these 19 crystals were compared to the hPrP-β2α2 structure solution from thin crystals discussed above. The correlation coefficient between the solved model and the individual crystals is shown in **Figure-2D** as CC_model_ (**Figure 3; SI Table 1**).

**Figure-3:**
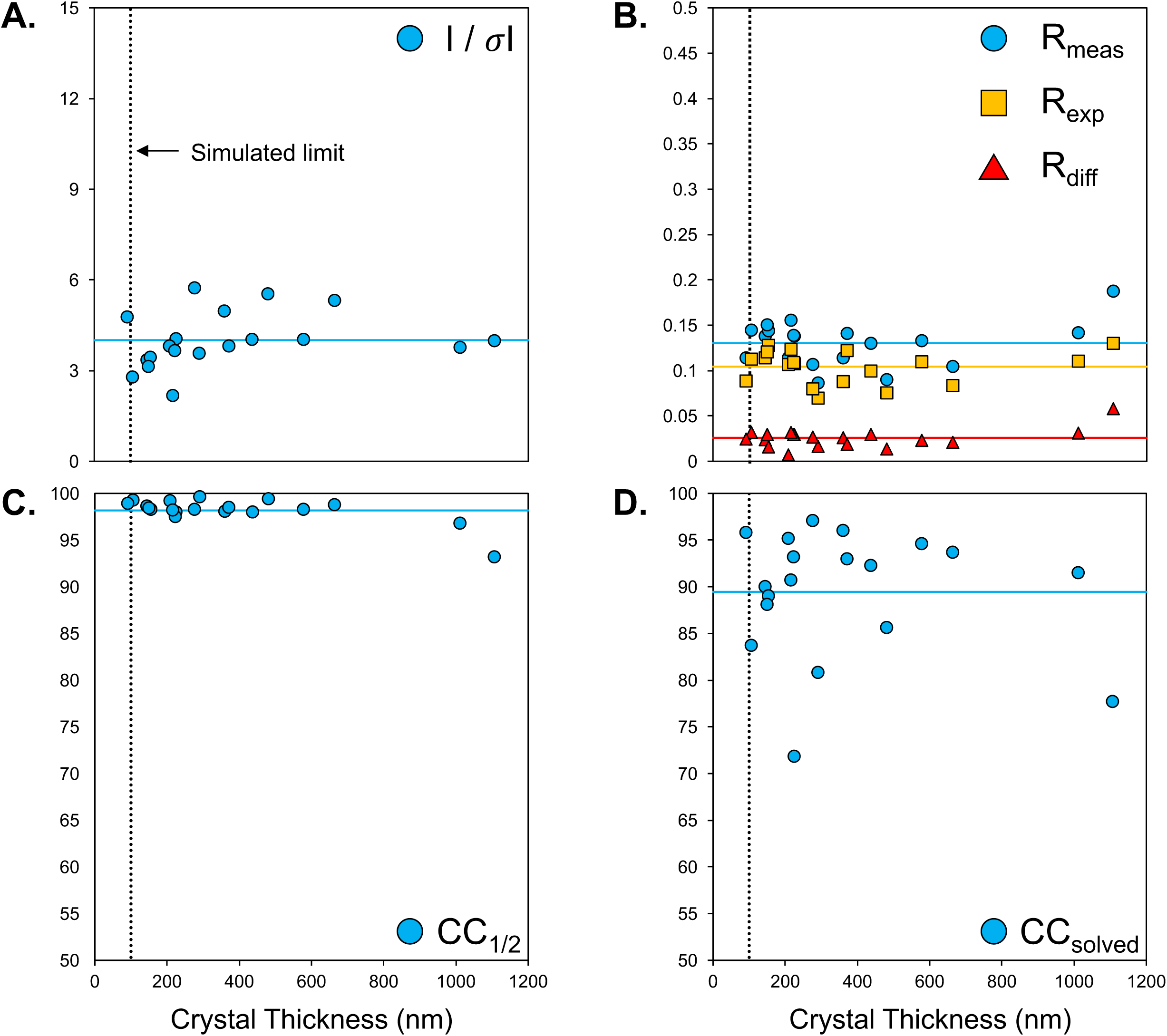
Measurements from hPrP-β2α2 crystals of variable thickness. (**A**) Mean I/ σI values, (**B**) Measured, expected R values and their difference, Rdiff, (**C**) internal half set correlation coefficient, and (**D**) the correlation of the measured structure factors to the solved thin crystal model, or CC_solved_. Mean values are depicted by solid lines. The dotted black line corresponds to the maximum reported limit from simulations (10).

#### Absorption by protein crystals

The data presented above indicates that dynamical scattering does not inhibit structure solution by MicroED even from micrometer thick crystals when data is collected by continuous rotation. However, we note that the achievable resolution was lower from thick crystals compared to thin crystals **(Figure 1,2 and Table 1)**. To evaluate whether absorption from thick crystals is limiting the achievable resolution, we recorded electron energy loss spectroscopy (EELS) spectra from 11 crystals of hPrP-β2α2 and 10 crystals of proteinase K with thicknesses ranging from ~100 nm to ~1400 nm at 300kV (**Figure 4**). Control spectra were recorded from regions of empty carbon support or within grid holes (**SI Document 3**). Each image was aligned by principal component analysis and a line scan through the zero-loss peak was measured along the first principal component. Intensities for the zero-loss peaks show expected exponential decay (9). A significant energy loss was observed for the carbon support alone with transmitted beam intensity decreasing by more than 40% (**SI Document 3**). An exponential fit to the data (**Figure 4**) suggests that the attenuation length, or (1/e) loss of intensity, due to a carbon film would be ~87 nm at 300kV. However, the (1/e) loss from crystals of proteinase K and hPrP-β2α2 are 261 nm and 323nm, respectively, at 300kV. Our estimates of the actual carbon thickness are about 45nm using intensity ratios (**SI Figure 6**), in good agreement with previous findings on these grids (24). Thus even after accounting for the carbon film support, our estimates are very close to the mean free path of water at 300kV, found experimentally to be 336nm (25, 26). This data suggests that even a three-fold decrease in intensity due to absorption is insufficient to prohibit structure solutions by MicroED for micrometer thick macromolecular crystals at 300kV; the achievable resolution drops quickly near or beyond this thickness limit.

**Figure-4:**
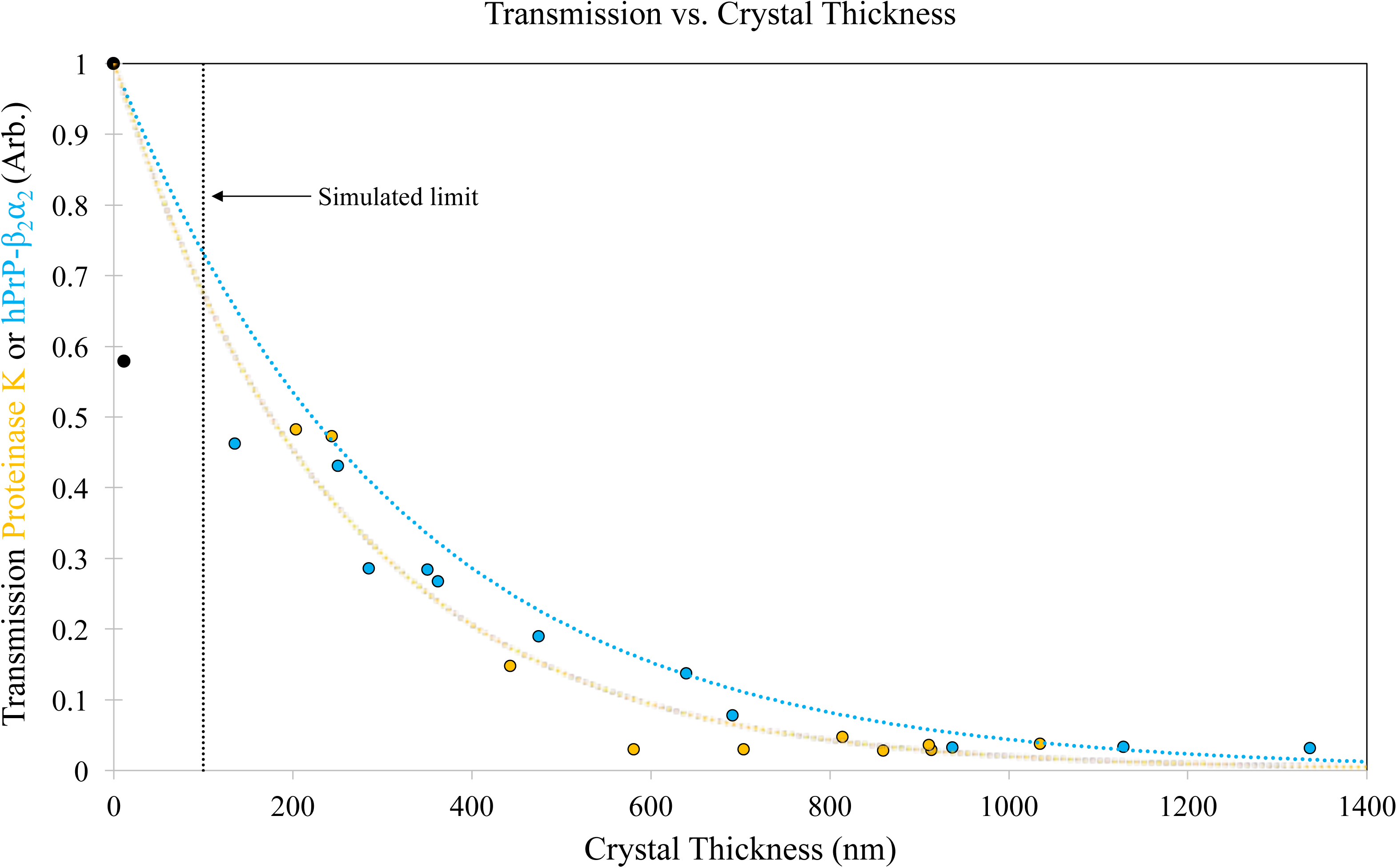
Observed transmission from crystals of various thicknesses. Peak intensities of the electron energy loss spectroscopy (EELS) spectra obtained from both hPrP-β2α2 (blue) and proteinase K (orange) crystals. Black dots correspond to the recorded vacuum intensity and the carbon film. Exponential decay models are presented as dashed lines coordinated to the sample color. The thickness limit for MicroED experiments suggested by simulatins is given as a dashed horizontal line (10).

## Discussion

These data indicate that accurate data can be collected and structures can be reliably determined by MicroED from crystals an order of magnitude thicker than previously expected. Simulations have suggested that structure determination by MicroED would be inhibited by dynamical scattering events from crystals only ~50-100nm thick (11, 27). Here we show that protein structure solutions can be obtained from substantially thicker crystals than simulated limits suggest for both globular proteins and peptides using both molecular replacement and *ab initio* methods, respectively.

High quality structure solutions can be obtained for both hPrP-β2α2 and proteinase K regardless of data originating from thin crystals or crystals nearly one micrometer thick. Clear atomic density in the 2F_o_-F_c_ map of hPrP-β2α2 reveals hydrogen atoms and the presence of zinc and acetate ligands as well as ordered water molecules. Multiple hydrogens are apparent in the F_o_-F_c_ density at the 3 σ level (**Figure 1**). Likewise, the density for proteinase K is of high quality with well-defined density in both the σ-weighted difference maps and the SA composite omit maps for data obtained from thin (<200nm), intermediate (~500nm) and thick (>1000nm) crystals. Thickness does not prevent data collection, integration, or structure determination from these crystals.

The initial structure solution for proteinase K determined by MicroED (**5i9s**) was refined to a resolution of 1.75 Å from merged data of four thin crystals (20). Our present models from single crystals of ~500nm and ~1000nm thickness are determined to 2 Å and 3Å resolution, respectively. We credit this difference in resolution to increased absorption as demonstrated by our EELS analyses from thicker crystals. Together these results indicate that the achievable resolution can be limited by thick specimens. However, merging data from multiple thick crystals yielded in improved statistics and therefore improved resolution, indicating that absorption phenomena can be overcome to some extent by increased redundancy in measurements. Thicker protein crystals are ultimately limited in resolution by absorption.

Why the discrepancy between the experimental data presented here and the limits found in previous simulations? The inherent difficulty of performing simulations requires assumptions of experimental conditions that fail to capture the complexity of MicroED experiments. Specifically, simulations assume that crystals are perfect, all the electrons are scattered and data is recorded from stationary crystals from major zone axes (11, 28). These conditions are often encountered in crystals of small molecules such as inorganic compounds in material science and from two-dimensional (2D) protein crystals, but are rarely encountered in macromolecular crystallography using 3D protein crystals and MicroED protocols (29). Crystals of macromolecules are highly mosaic compared with those of simple organic compounds or inorganic crystals. Macromolecular crystals are imperfect and bent at the nanoscale. MicroED data from highly mosaic crystals benefits from their disorder in analogy to the pivoting of an electron beam in precession electron diffraction. Moreover, MicroED data is collected by continuous rotation so integration over the rocking curve further curbs dynamical effects. Our results confirm that dynamic scattering is not a major problem in solving protein crystal structures, and that dynamic scattering effects do not increase linearly with crystal thickness. However, the question of why significant artifacts are not observed from multiple scattering still remains, and requires further study.

The fact that multiple scattering artifacts do not limit structure determination in MicroED is especially surprising for crystals thicker than ~300nm, where the crystalline thickness easily exceeds the mean free path of an electron at either 200 or 300kV (**Figure 4**) (25, 26, 30). For these crystals, few electrons are transmitted through the crystal without any interaction with the specimen. At very high thicknesses the diffraction spots begin to widen due to energy loss. Larger spots make crystals with many atoms per unit cell difficult to measure and may give poor statistics as a large fraction of scattering will be reduced by absorption. Integrating large broadened spots from crystals with large unit cells may result in spot-overlap and other complications that may limit the achievable resolution; we have yet to encounter this problem. The largest protein determined to date by MicroED is catalase at ~240kDa and even with such a large unit cell and peak broadening, spot overlap was not observed and did not hinder structure determination (14). While broadening of diffraction spots can be curbed using an energy filter, this comes at the cost of reducing the transmitted signal. Nonetheless, use of an energy filter could improve signal to noise in MicroED experiments and the influence of thick crystals with large unit cells on diffraction quality remains to be further investigated.

In determining protein structures from crystals up to 1 micrometer in thickness using both *ab initio* phase retrieval and molecular replacement, we demonstrate that MicroED experiments are ultimately limited by absorption effects and crystal quality. At present, we estimate crystals ~500nm or thinner maximize data quality (~2x mean free path for 200kv), but structures from thicker crystals are not necessarily precluded. In fact, our results from even thicker crystals provides confidence that observed diffraction can be free of multiple scattering artifacts in a typical continuous rotation MicroED experiment, even from micrometer thick protein crystals.

## Concluding Remarks

We demonstrate that MicroED yields accurate diffraction intensities, and that structures can be determined by molecular replacement or *ab initio* methods from protein crystals much thicker than previously suggested. Our data demonstrates that, as with X-ray diffraction, dynamical scattering is not a prohibitive source of error when MicroED data is collected on protein crystals by continuous rotation. Instead, absorption phenomena limit the achievable resolution in a way that can be curbed by merging multiple data sets. We suggest a stringent upper bound on crystal thickness of about one micrometer on MicroED experiments and a soft upper bound of about 500nm to ensure high-quality diffraction with the best possible resolution; these limits are not imposed by dynamical scattering, but instead by absorption. Our study expands the usefulness of MicroED as a general method for structure determination to atomic resolution from specimens up to a micrometer thick, opening new avenues of research that may broadly impact structural biology.

## Materials and Methods

### Protein preparation

hPrP-β2α2 with greater than 98% purity was purchased from Genscript, dissolved in water at 10-20mg/ml and screened by the hanging drop method in a high throughput screen. Initial hits were optimized in 24-well hanging drop trays. The best crystals were observed in a condition containing 10% (w/v) PEG-8000; 0.1 M MES pH 6.0; Zn(OAc)2. This condition was used as the basis for a batch crystallization of the peptide at 10 mg/ml at a 1:1 ratio of peptide solution to mother liquor. In this condition, crystals grew as needle clusters that could be broken by force of pipetting and applied to grids for cryopreservation. Proteinase K (*E. Album*) was purchased from Sigma and used without further purification. Crystals were grown by adding 5 μL of protein solution (50 mg/ml) to 5 μL of precipitant solution (1.5 M ammonium sulfate, 0.1M Tris pH 8.0) in a sitting drop vapor diffusion tray. Large proteinase crystals were collected from sitting drops and sonicated into smaller crystals as previously described (4).

### MicroED data collection

MicroED data was collected for 19 crystals of the peptide hPrP-β2α2 (Gly-Ser-Asn-Gln-Asn-Asn-Phe) of varying thickness over the real space wedge corresponding -30° to +30° to under continuous rotation. All MicroED experiments were performed on an FEI Tecnai F20 microscope at an accelerating voltage of 200 kV, corresponding to a wavelength of 0.0251 Å. Data was collected using a constant rotation rate of 0.2° s^-1^ on a TVIPS TemCam-F416 CMOS detector in rolling-shutter mode with 5 s exposures. Beam intensity for all hPrP-β2α2 crystals was constant with an average dose rate of 0.003e^-^ Å^-1^ sec^-1^, or an overall exposure of ~1 e^-^Å^-2^. Proteinase K crystals were collected at a dose rate of 0.01 e^-^ Å^-1^ sec^-1^ with an overall total dose per crystal of less than 3 e^-^ Å^-2^. hPrP-β2α2 samples maintained a camera length of 0.730 m equivalent to a sample–detector distance of 1.313 m in a corresponding lens less system. Proteinase K data were collected at a camera length of 1.2m, corresponding to a detector distance of 2.200m. Diffraction data were collected through a circular selected area aperture of 1 μm^2^ in projection to reduce background noise. All TEM measurements were done at liquid nitrogen temperatures (~77 K).

### Determination of Crystal Thickness

Crystal thickness was assessed in two ways: geometrical measurement and by using counts on the camera. First, Average crystal thickness was initially estimated geometrically by examining the projected area of each crystal from images taken at 0°, 15°, 30°, 45°, and 60° using the pixel length calibration from the known grid hole size of either 1 or 2 μm in ImageJ (NIH). Crystals of hPrP-β2α2 were all rod-shaped and assumed to be ellipsoidal rods. Tilt series assessed the aspect ratio of the major (**a**) and minor (**b**) axis at each angle, with a known length of 2**a** being the rod width measured at 0°. The average thickness of a crystal is then π/4 = ~0.785% of the maximum thickness, 2**b**, perpendicular to the film. Proteinase K crystals were assumed to be cuboids with edge lengths **w**, **l**, **h**. Edge lengths of **w** and **l** were found from the 0° images and the **h** edge was estimated by the change in projected area of the crystal over a fixed length along the **x** and **y** axis. The average thickness of these crystals was assumed to be equal to the estimation of the **h** edge. Measurement accuracy was between 5 and 10 nm, the average variance for each crystal was ~20%, and the standard deviation between intensity and geometry estimations was ~100nm. Second, The geometrical approximation of crystal thickness above was validated by using intensity ratios as previously described (2, 14, 26). Each crystal from the hPrP-β2α2 set in **Figure-3**/**SI Document 2** at 0° were identified at 0°. Each crystal had an area selected from the portion of the crystal collected upon where the median intensity transmitted was calculated. The same area was then used to calculate the median intensity for the nearest clean carbon area and nearest empty grid hole for comparison. The crystal thickness was estimated by first estimating the carbon thickness using the formula given in Feja & Aebi (26) assuming all missing intensity was lost due to energy loss signals and the incident maximum intensity was that of the beam intensity in a grid hole. The mean free path of the amorphous carbon was calculated using the average energy loss of carbon being 14.1 eV derived from the formula given in (26), with Z=6, corresponding to a mean free path of 125nm at 200kV. This gave a consistent value of the carbon thickness being approximately 45nm with a standard deviation of 7nm. The crystal thickness is then calculated by subtracting the contribution of the carbon using the same equation with a mean free path of 242nm, calculated by scaling the value of 203nm mean free path of vitreous ice given by Grimm et al. at 120kV (25, 30). Our two measures were found to have a standard deviation of approximately 100nm. The correlation coefficient between the two measures was found to be 89%. This suggests our geometrical measurements are accurate within 25% of our listed values. The correlation plot and deviations from these intensity measures are given in **SI Figure 6**.

### Absorption Experiments

Energy spectra were collected on a JEOL JEM-3200FSC microscope at 300 kV with a 500 nm^2^ aperture in projection and dose of 0.1 e^-^ total Å^-2^. Spectra were collected on a TVIPS TemCam-F416 CMOS detector at full resolution in normal (integration) mode and saved as 16-bit signed integer TIFF files. Spectra were scaled in ImageJ to the known 100 eV μm^-1^ energy filter spacing and 15.6 μm pixel size. Individual images were loaded into Mathematica and aligned with their principal component along the horizontal axis – typically resulting in a 6.7° clockwise rotation. Line scans of width 1 pixel were selected through the zero-loss peak along the principal loss axis, and shifted such that their peak intensities were located at an energy loss of 0 eV. Thickness for crystals used in spectra were determined as described above. Zero-loss peaks were fit to a general Gaussian model of I=a*e^-b(x-c)2^ to determine peak maxima and integrated peak intensity. Data points were weighted by the square root of their intensities for the fitting to assure proper solutions. Zero-loss peak intensity was fit to a general exponential decay model with the intercept fixed to the normalized value of the vacuum peak as in **Figure-4**. Exponential fits to the experimental data for the carbon film, proteinase K, and hPrP-β2α2 protein crystals resulted in R^2^ residuals of 1.0, 0.85, and 0.88, respectively.

### MicroED Data Processing

Diffraction movies were converted to the SMV file format using TVIPS tools (31) and checked for pixel truncation as previously described (21). Indexing and integration were performed in XDS (20, 32). Integrated diffraction intensities were sorted and merged in XSCALE. Merged intensities were converted to amplitudes and the files formatted to SHELX format in XDSCONV. Thick and thin hPrP-β2α2 were solved using SHELXT, placing all of the atoms in the unit cell correctly (33). SHELXT supported the hypothesis of P1 crystallographic symmetry (16). Intensities for proteinase K were input directly from XSCALE into Phaser for molecular replacement (34). Molecular replacement was successful with a LLG > 1000 and a TFZ > 20 using the model **5i9s**. Refinement was carried out using phenix.refine (22). Individual models were adjusted manually in Coot (35) by visual inspection of the atomic model against the F_o_-F_c_ and 2F_o_-F_c_ maps. All-atom composite omit maps with simulated annealing were calculated in Phenix (22).

### Models and Figures

Figures were generated in either PyMol or VMD (36, 37). Plots and fits to data such as EELS spectra were created in Mathematica using nonlinear model fits to the data. Backbone RMSDs were calculated in VMD. All atom RMSD values were generated by the align command in PyMol.

## Acknowledgements

We thank David Eisenberg (HHMI, UCLA), Robert Glaeser (UCB), and Michael Sawaya (UCLA) for helpful discussions and/or critical reading of this manuscript. Jose Rodriguez is supported as a Searle Scholar and a Beckman Young Investigator. The Gonen Laboratory is supported by the Howard Hughes Medical Institute. This work was also supported by the Janelia Research Visitor Program.

## Table and Figure Legends

**SI-Table-1:** MicroED diffraction data from hPrP-β2α2 crystals of various thicknesses.

| Mean Thickness (nm) | # Observed | # Unique | # Possible | Completeness | R<sub>observed</sub> | R<sub>expected</sub> | # Compared | I/σI | R<sub>measured</sub> | CC<sub>1/2</sub> | CC<sub>model</sub> | R<sub>difference</sub> | Crystal # |
| --- | --- | --- | --- | --- | --- | --- | --- | --- | --- | --- | --- | --- | --- |
| 91 | 1420 | 771 | 2625 | 0.29 | 0.08 | 0.09 | 1298.00 | 4.78 | 0.11 | 98.90 | 95.80 | 0.03 | 18 |
| 106 | 921 | 518 | 2490 | 0.21 | 0.10 | 0.11 | 806.00 | 2.79 | 0.15 | 99.30 | 83.70 | 0.03 | 11 |
| 145 | 1311 | 720 | 2391 | 0.30 | 0.10 | 0.11 | 1182.00 | 3.37 | 0.14 | 98.60 | 90.00 | 0.02 | 7 |
| 150 | 1031 | 584 | 2622 | 0.22 | 0.11 | 0.12 | 894.00 | 3.14 | 0.15 | 98.40 | 88.10 | 0.03 | 20 |
| 155 | 1266 | 676 | 2544 | 0.27 | 0.10 | 0.13 | 1180.00 | 3.46 | 0.14 | 98.30 | 89.00 | 0.02 | 12 |
| 209 | 1390 | 747 | 2596 | 0.29 | 0.08 | 0.11 | 1286.00 | 3.82 | 0.11 | 99.20 | 95.20 | 0.01 | 5 |
| 217 | 1313 | 700 | 2582 | 0.27 | 0.11 | 0.12 | 1226.00 | 2.19 | 0.16 | 98.20 | 90.70 | 0.03 | 13 |
| 223 | 1396 | 767 | 2566 | 0.30 | 0.10 | 0.11 | 1258.00 | 3.67 | 0.14 | 97.50 | 93.20 | 0.03 | 8 |
| 225 | 1280 | 690 | 2587 | 0.27 | 0.10 | 0.11 | 1180.00 | 4.07 | 0.14 | 98.00 | 71.80 | 0.03 | 3 |
| 277 | 1415 | 775 | 2585 | 0.30 | 0.08 | 0.08 | 1280.00 | 5.74 | 0.11 | 98.30 | 97.10 | 0.03 | 21 |
| 291 | 942 | 572 | 2543 | 0.23 | 0.06 | 0.07 | 740.00 | 3.59 | 0.09 | 99.60 | 80.80 | 0.02 | 14 |
| 360 | 1471 | 790 | 2624 | 0.30 | 0.08 | 0.09 | 1362.00 | 4.97 | 0.11 | 98.10 | 96.00 | 0.03 | 9 |
| 372 | 1419 | 769 | 2526 | 0.30 | 0.10 | 0.12 | 1300.00 | 3.83 | 0.14 | 98.50 | 93.00 | 0.02 | 17 |
| 437 | 1351 | 725 | 2526 | 0.29 | 0.09 | 0.10 | 1252.00 | 4.04 | 0.13 | 98.00 | 92.30 | 0.03 | 10 |
| 481 | 1280 | 689 | 2665 | 0.26 | 0.06 | 0.08 | 1182.00 | 5.54 | 0.09 | 99.40 | 85.60 | 0.01 | 1 |
| 578 | 1505 | 816 | 2644 | 0.31 | 0.09 | 0.11 | 1378.00 | 4.05 | 0.13 | 98.30 | 94.60 | 0.02 | 16 |
| 664 | 1504 | 796 | 2554 | 0.31 | 0.07 | 0.08 | 1416.00 | 5.34 | 0.11 | 98.80 | 93.70 | 0.02 | 2 |
| 1012 | 1157 | 597 | 2593 | 0.23 | 0.10 | 0.11 | 1060.00 | 3.77 | 0.14 | 96.80 | 91.50 | 0.03 | 15 |
| 1107 | 686 | 378 | 2180 | 0.17 | 0.13 | 0.13 | 616.00 | 3.99 | 0.19 | 93.20 | 77.70 | 0.06 | 19 |

**SI-Figure-1:**
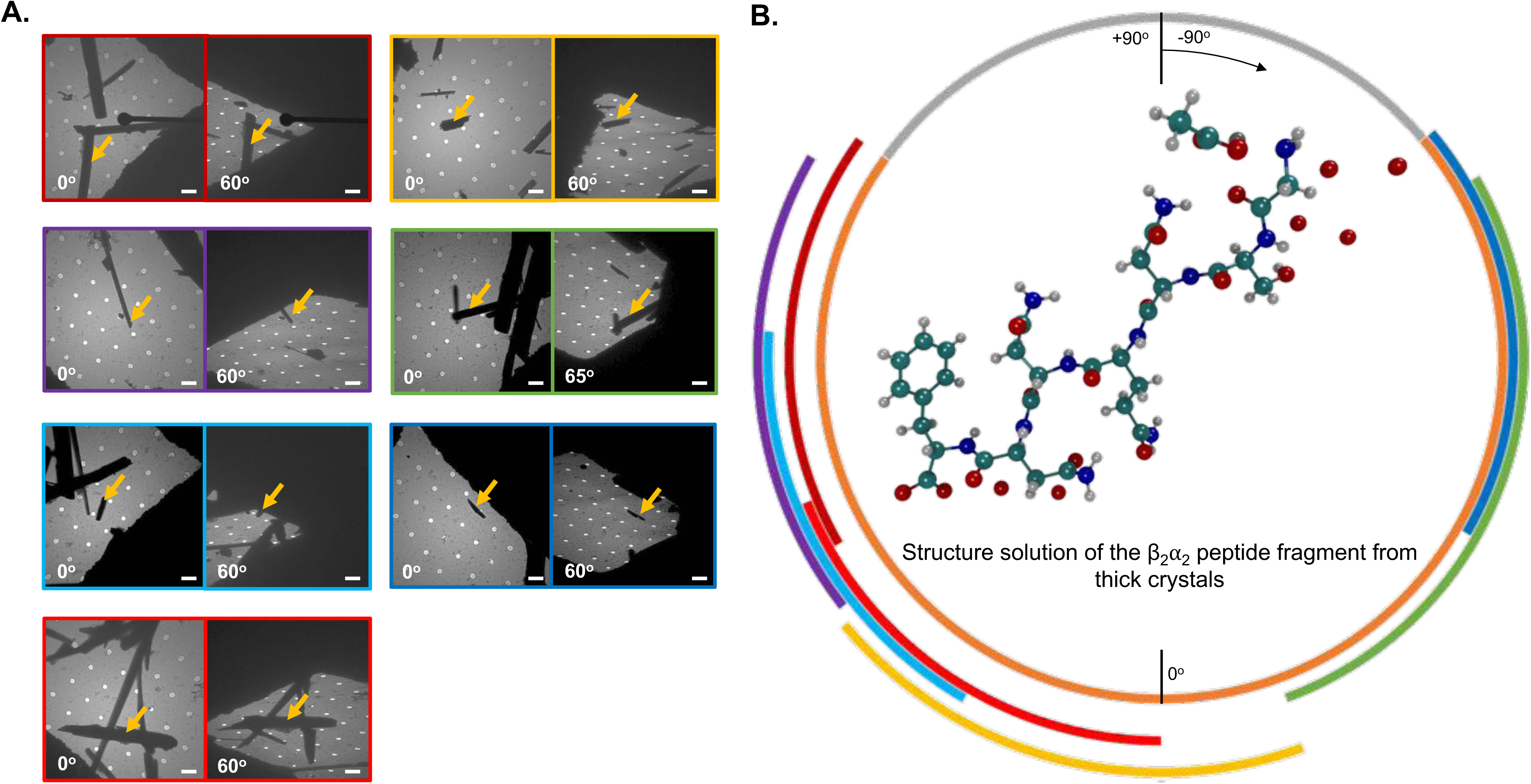
(**A**) Thick crystals used to solve the thick hPrP-β2α2 structure. (**B**) Structure solution of hPrP-β2α2 solved from 7 crystals with average thickness >1 μm. Colors show individually measured angular ranges between -65 and -63 degrees with corresponding crystals having matching border colors. Grey regions correspond to the angular wedge not measured and (orange) showing the region available for measurement. Scale bars correspond to 2 μm.

**SI-Figure-2:**
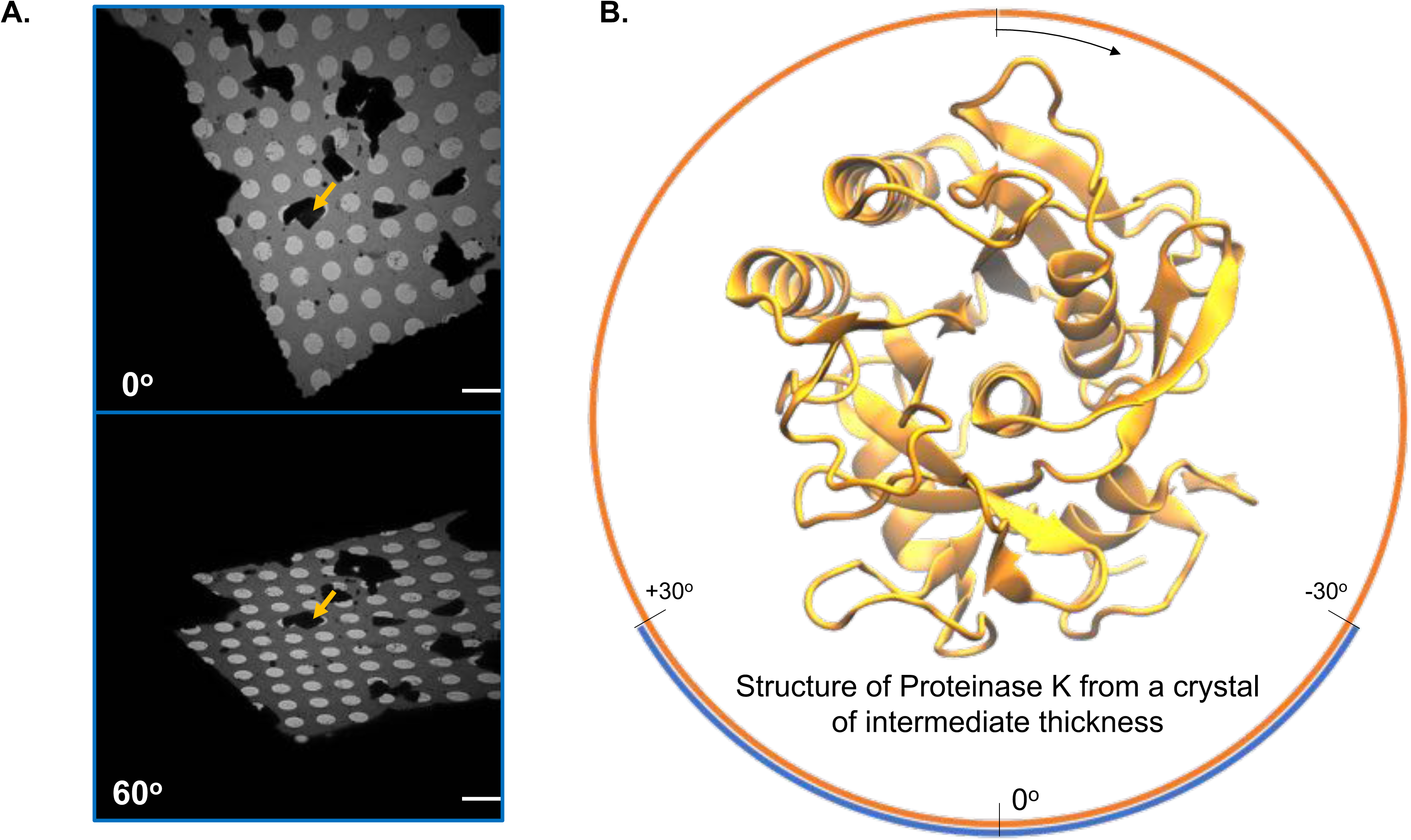
(**A**) Single 500 nm crystal used to solve the proteinase K structure at marked angles and locations. (**B**) Corresponding structure solution model. Measurable data after inclusion of symmetry (orange) and recorded angular range (blue) from the crystal. Grey angular regions depict ranges not measured. Scale bars correspond to 2 μm.

**SI-Figure-3:**
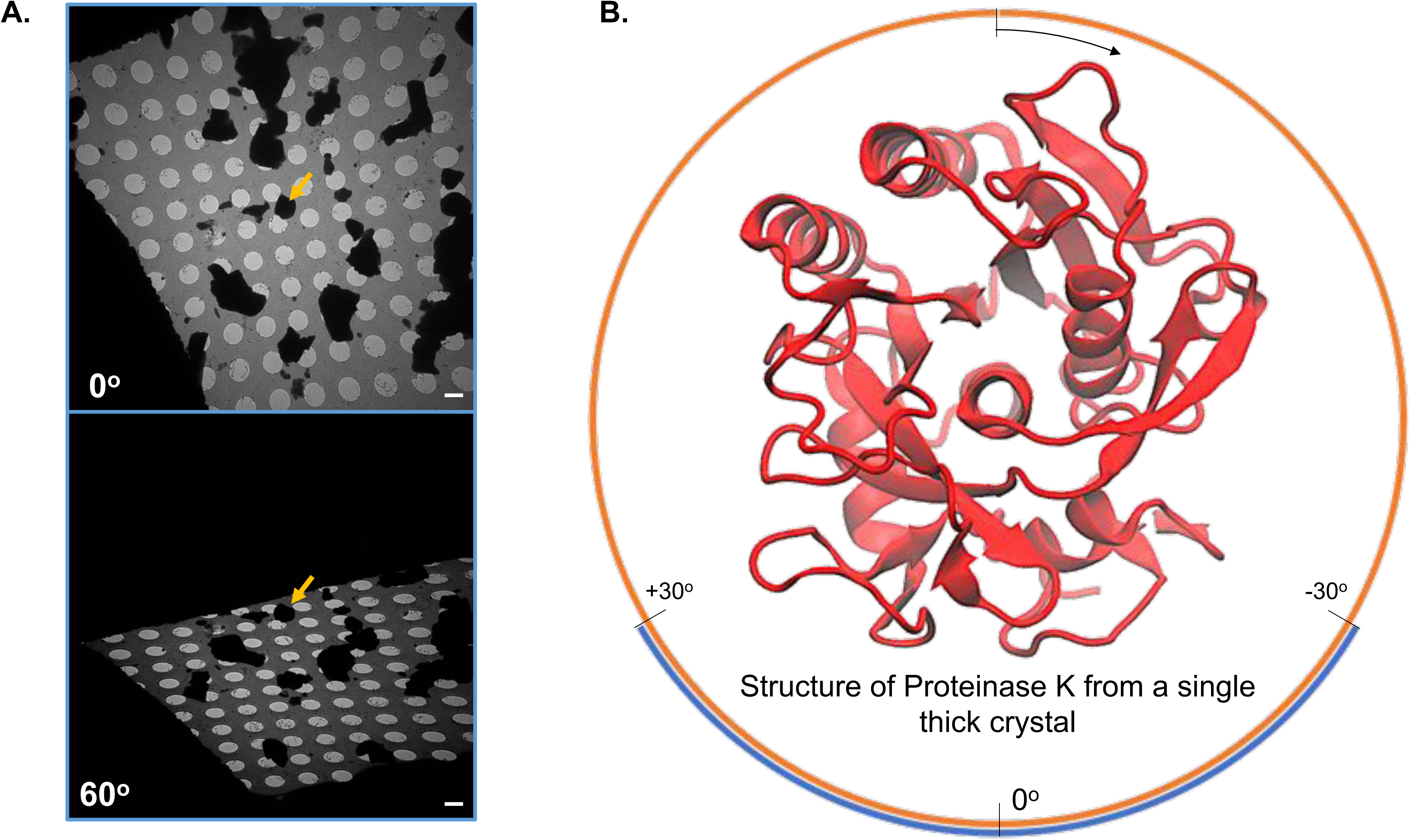
(**A**) Single 1 μm crystal used to solve the proteinase K structure at marked angles and locations. (**B**) Corresponding structure solution model. Measurable data after inclusion of symmetry (orange) and recorded angular range (blue) from the crystal. Grey angular regions depict ranges not measured. Scale bars correspond to 2 μm.

**SI-Figure-4:**
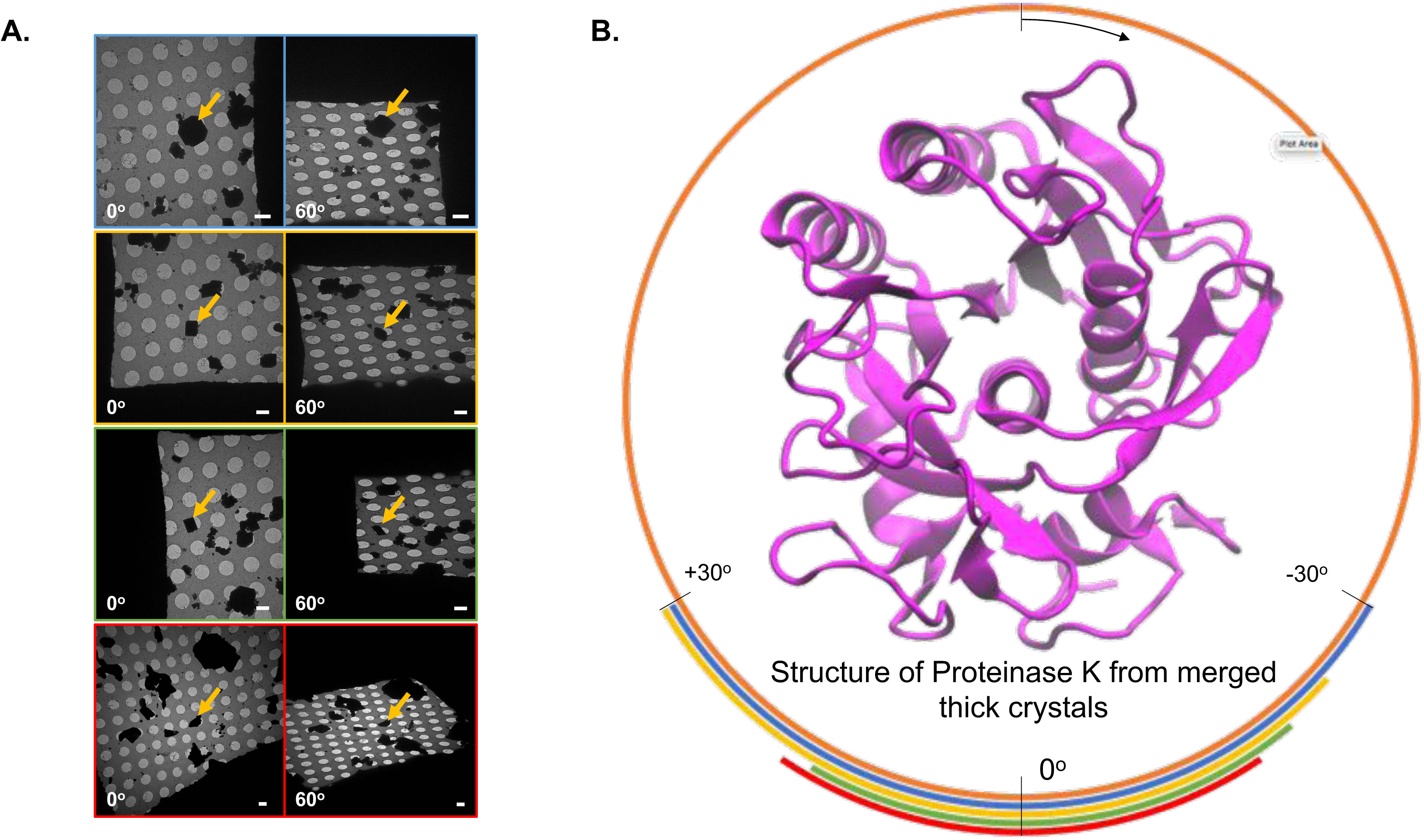
(**A**) Four >1 μm crystals used to solve the proteinase K structure at marked angles and locations. (**B**) Corresponding structure solution model. Measurable data after inclusion of symmetry (orange) and recorded angular range (blue) from the crystal. Grey angular regions depict ranges not measured. Scale bars correspond to 2 μm.

**SI-Figure-5:**
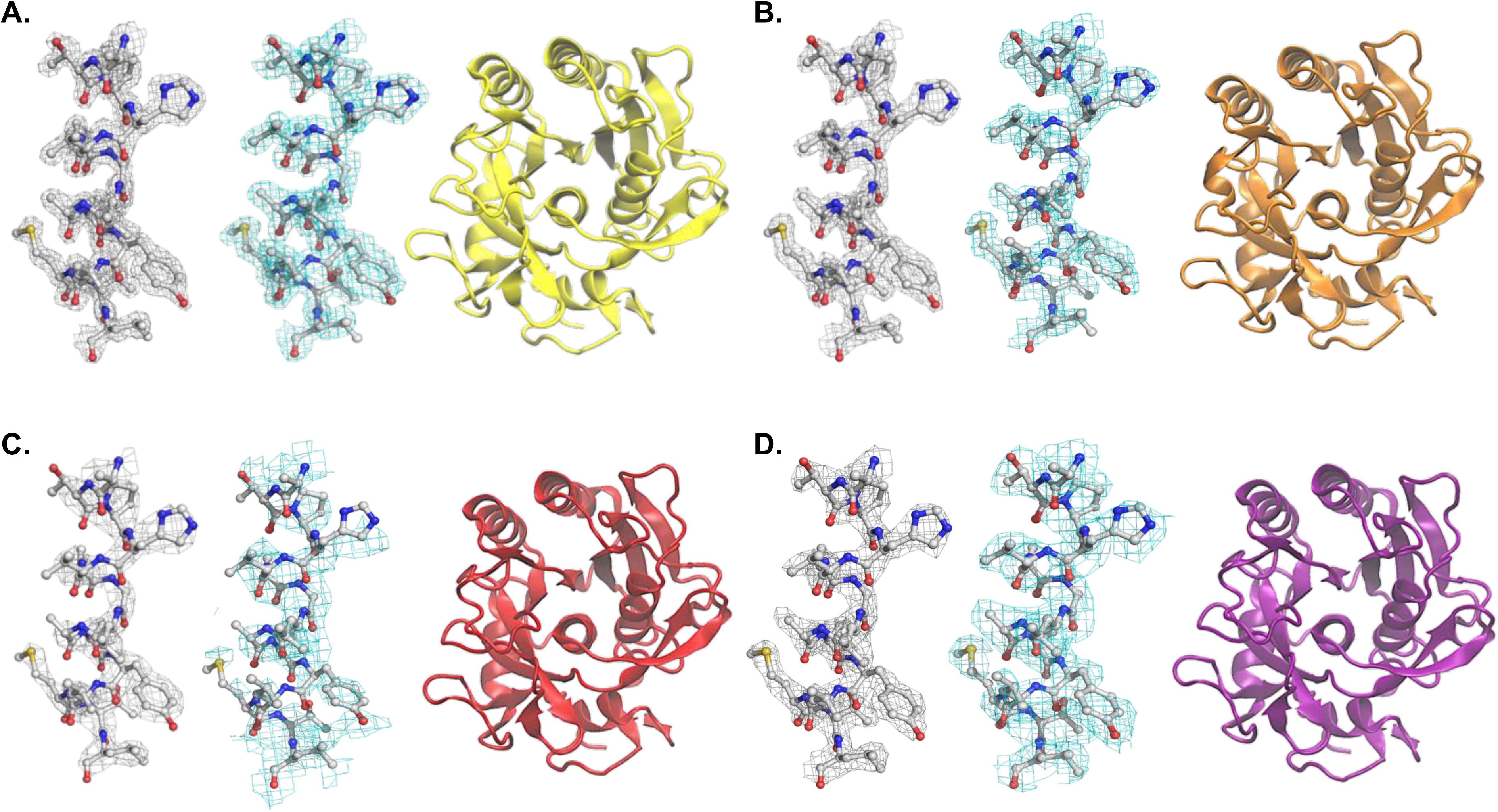
2mFo-DFc and all-atom SA composite omit potential maps with corresponding structural model for proteinase K from (**A**) four thin crystals, (**B**) a single 600 nm, (**C**) a single 1 μm, and (**D**) four merged crystals >1 μm thick. 2F_o_-F_c_ potential maps are contoured at the 1.5σ level with a 2 Å curve.

**SI-Figure-6:**
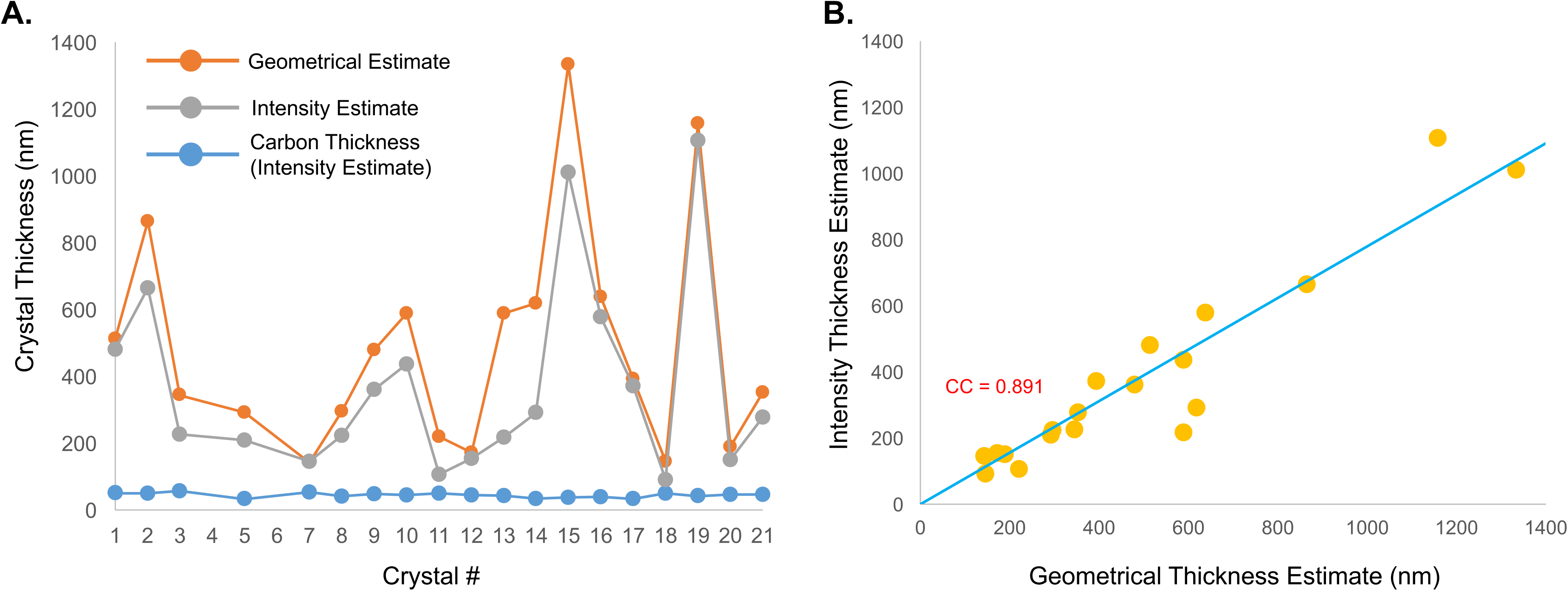
Crystal thickness estimation. (**A**) Comparison between the geometrically measured thickness (orange) and the thickness estimate from the Beer-Lambert Law (Grey), and the calculated carbon thickness from the Beer-Lambert calculation (Blue). (**B**) Correlation between the two types of measures of crystal thickness.

**SI-Document-2:** Peptide tilt angle images (diffraction data crystals) - PDF

**SI-Document-3:** EELS data (crystals, spectra, and fits) - PDF

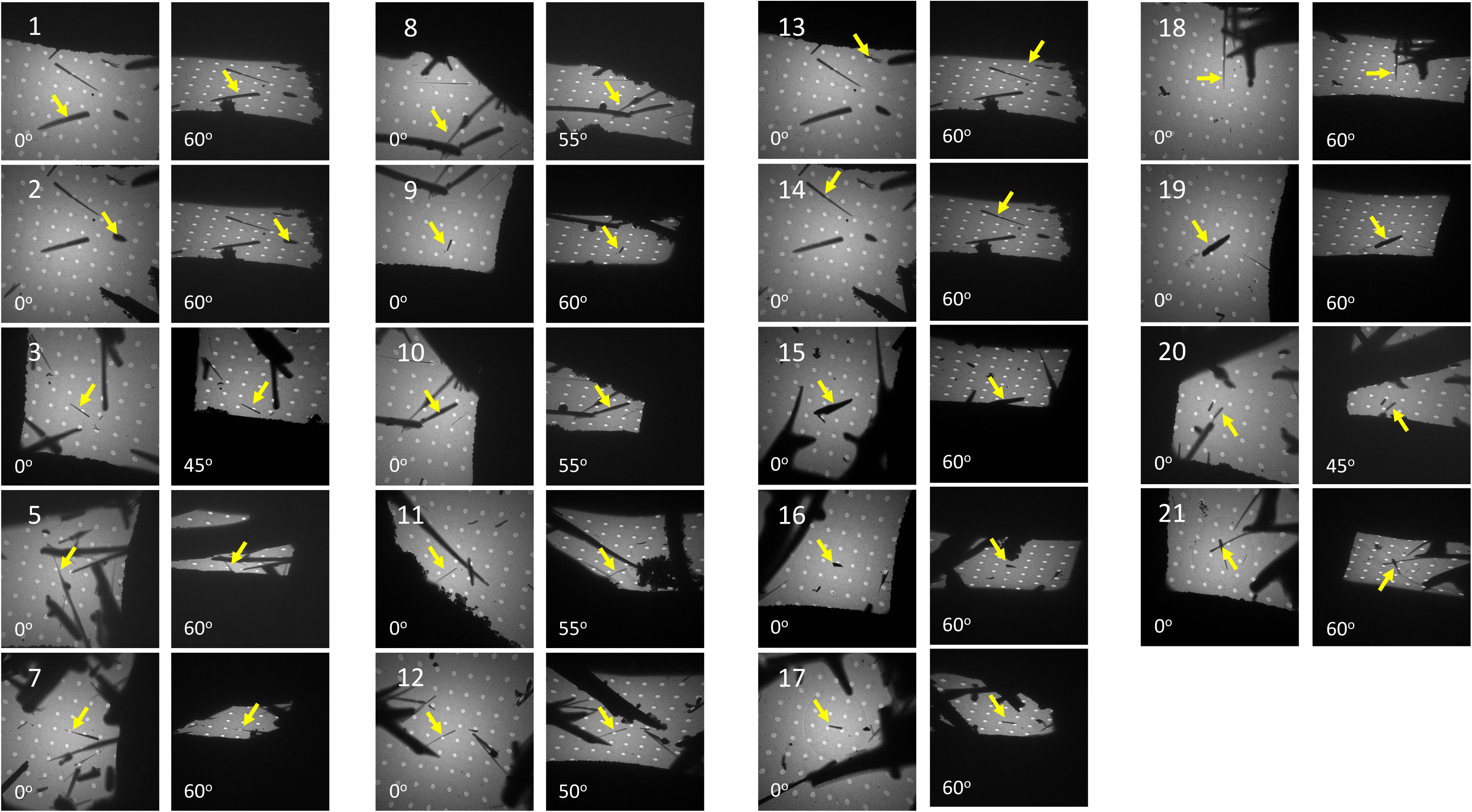
Crystals of hPrP - β_2_α_2_ used to generate statistics in Figure-3 and SI-Table-1

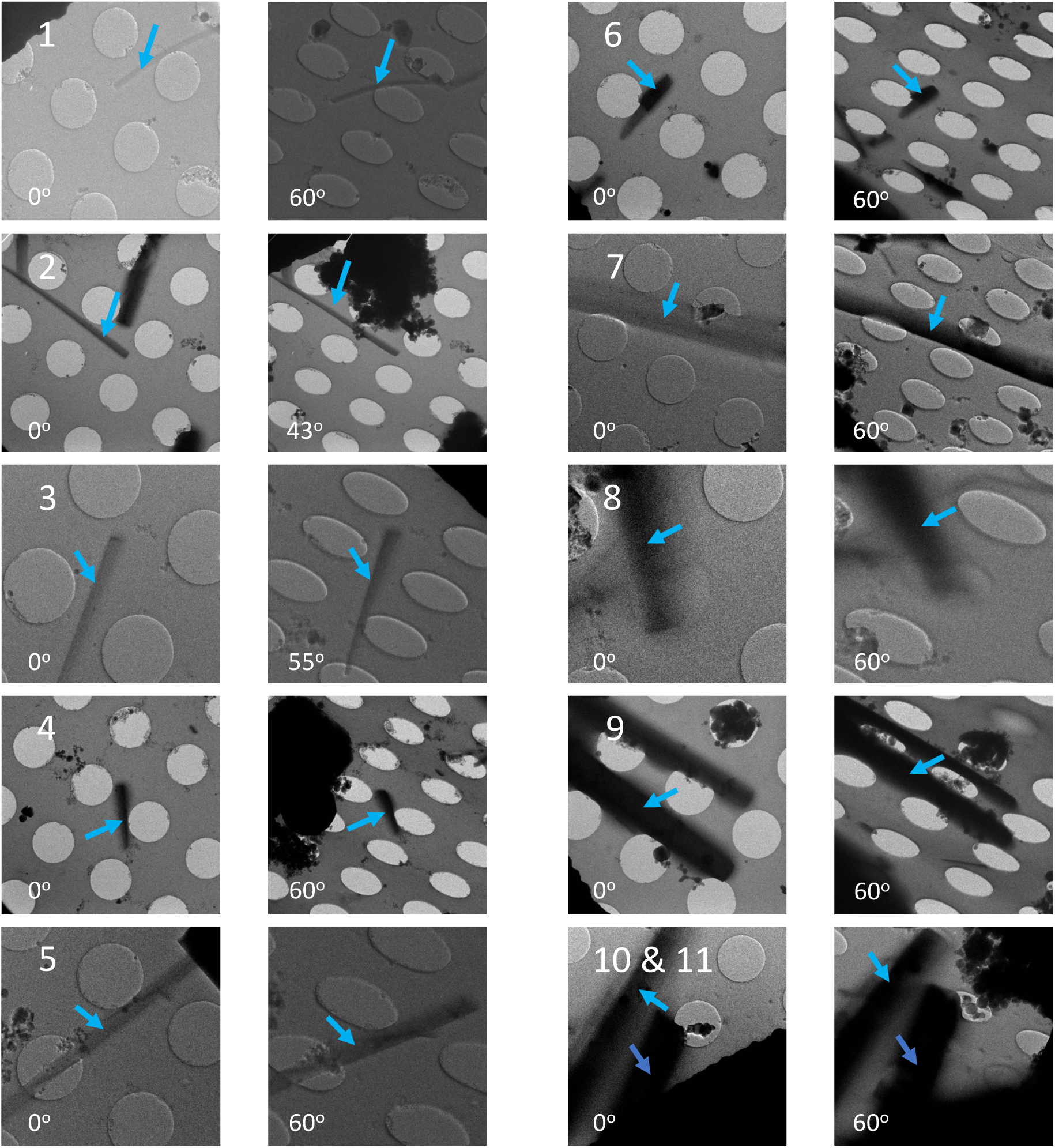
hPrP - B_2_α_2_ Crystals used in EELS Experiments

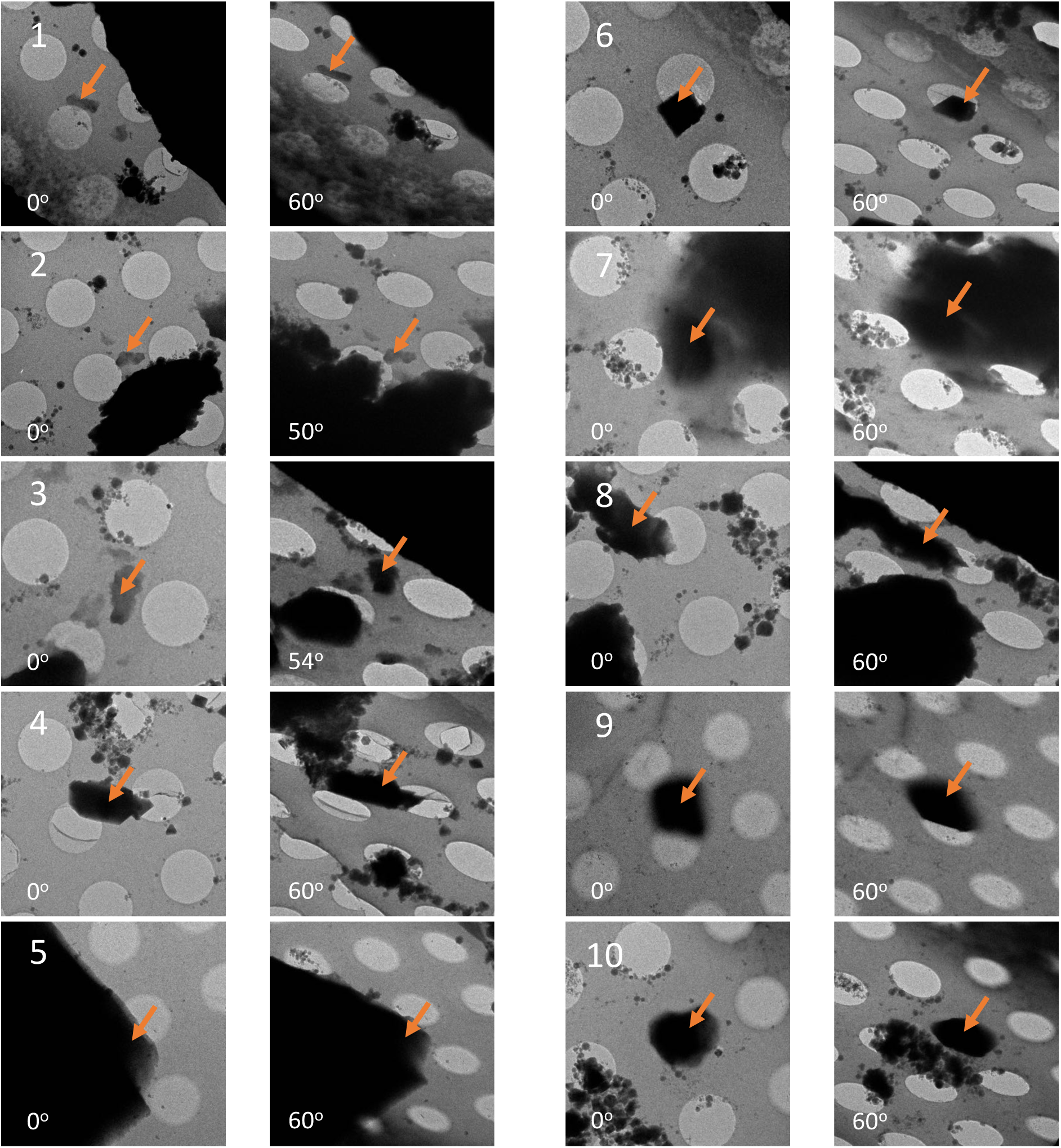
Proteinase K Crystals in EELS Experiments

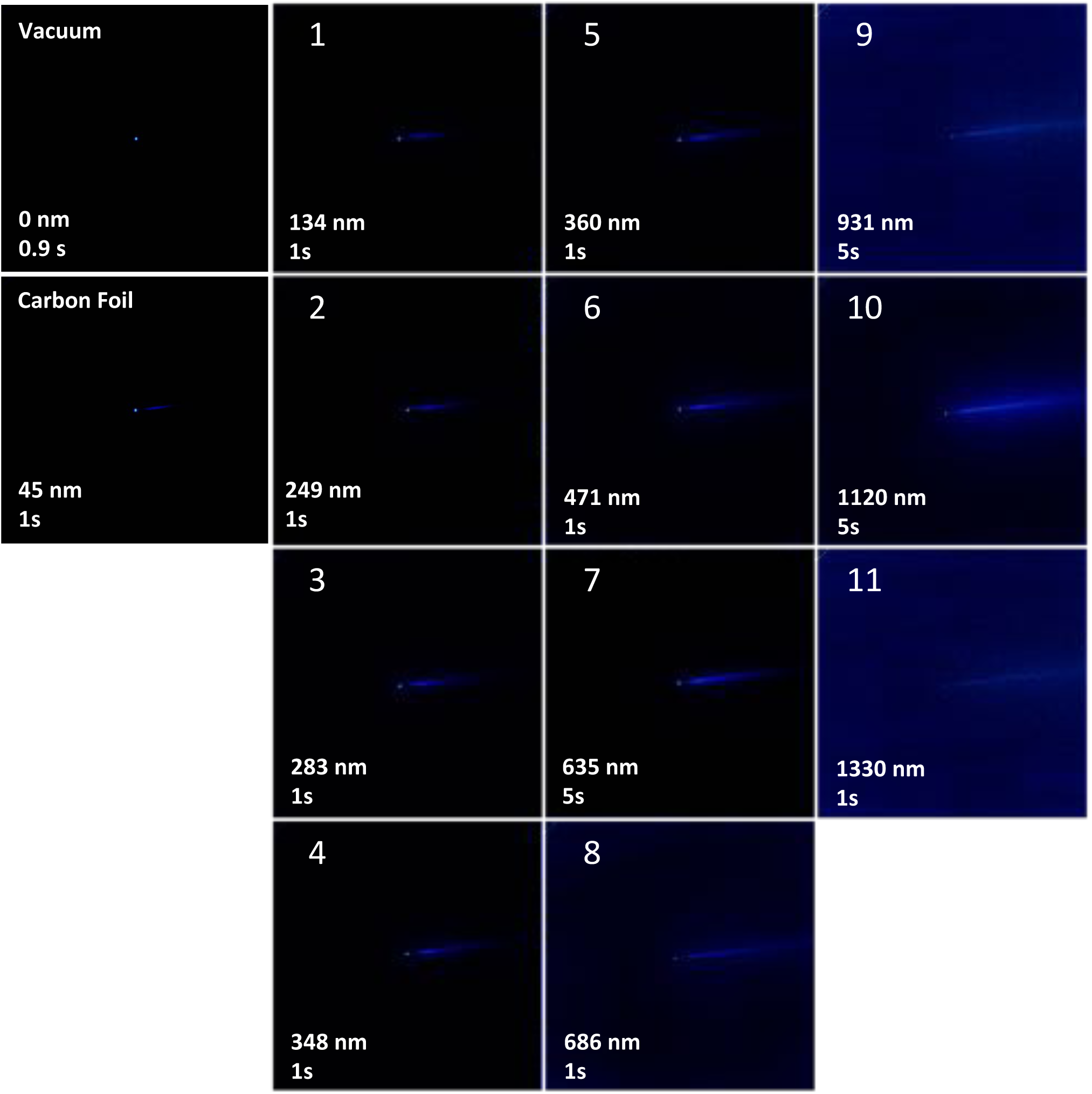
EELS Spectra from β_2_α_2_ crystals at 300 kV

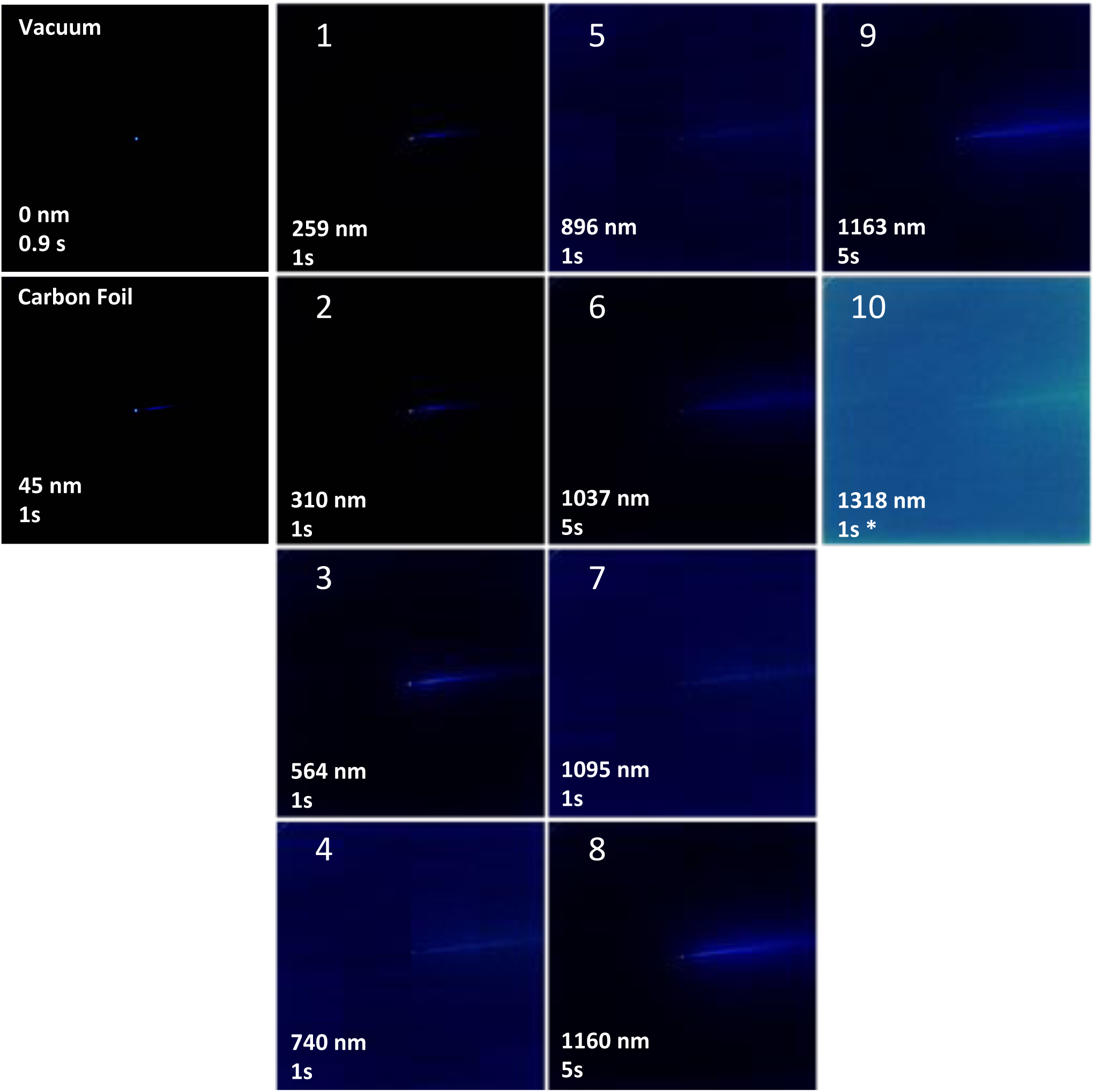
EELS Spectra from Proteinase K Crystals at 300kV

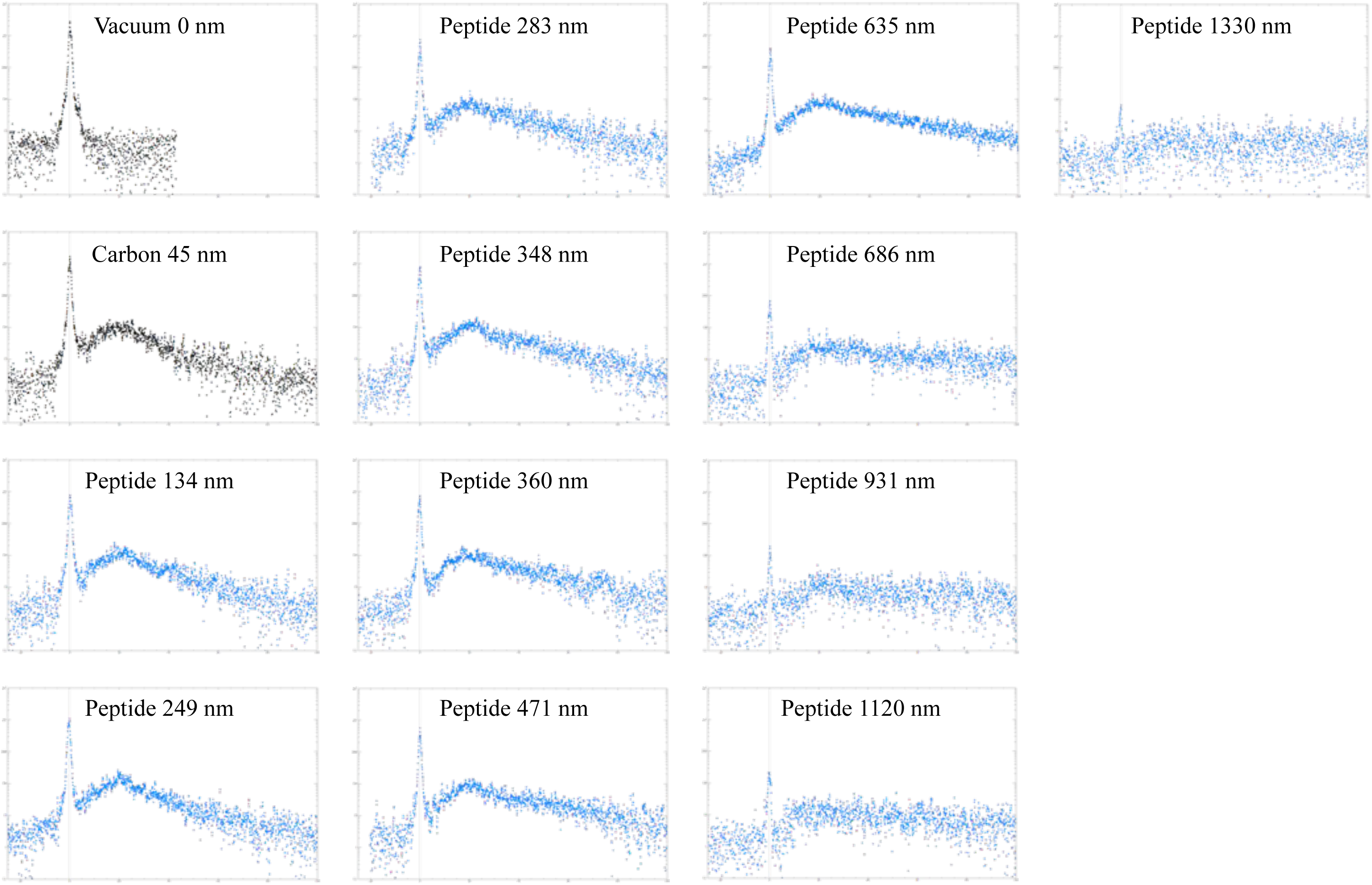
Line Scans of EELS Spectra for hPrP - β_2_α_2_ at 300 kV

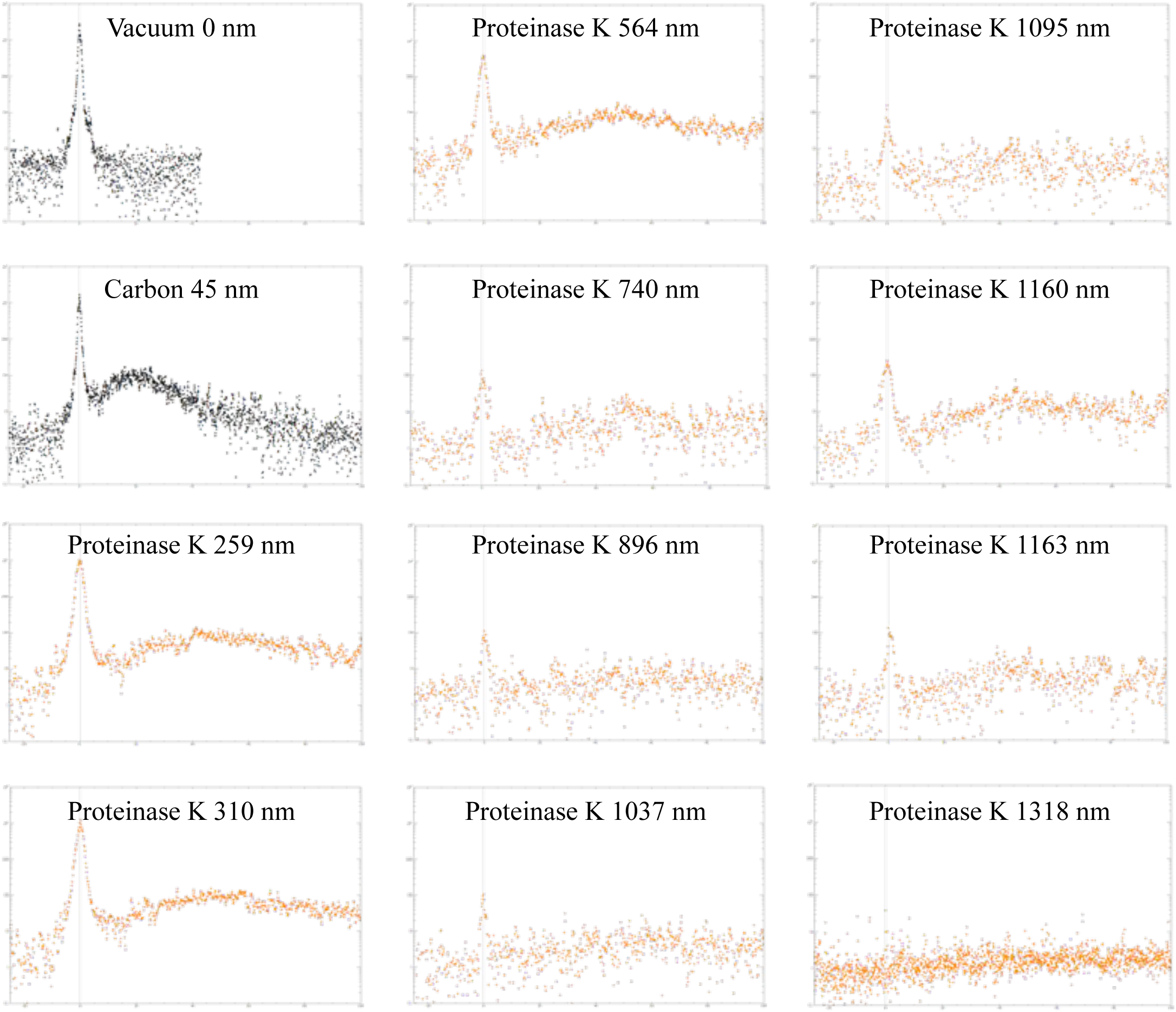
Line Scans of EELS Spectra for Proteinase K Crystals

